# Resting-state fMRI-based screening of deschloroclozapine in rhesus macaques predicts dosage-dependent behavioral effects

**DOI:** 10.1101/2021.11.24.469738

**Authors:** Atsushi Fujimoto, Catherine Elorette, J. Megan Fredericks, Satoka H. Fujimoto, Lazar Fleysher, Peter H. Rudebeck, Brian E. Russ

**Author notes:** These authors contributed equally to this work (Atsushi Fujimoto, Catherine Elorette). Joint last author (Peter H. Rudebeck, Brian E. Russ). Corresponding author, **Correspondence:** Brian E. Russ.

## Abstract

Chemogenetic techniques such as Designer Receptors Exclusively Activated by Designer Drugs (DREADDs) enable transient, reversible, and minimally invasive manipulation of neural activity in vivo. Their development in non-human primates is essential for uncovering neural circuits contributing to cognitive functions and their translation to humans. One key issue that has delayed the development of chemogenetic techniques in primates is the lack of an accessible drug-screening method. Here, we utilize resting-state functional MRI (rs-fMRI), a non-invasive neuroimaging tool, to assess the impact of deschloroclozapine (DCZ) on brain-wide resting-state functional connectivity in seven rhesus macaques without DREADDs. We found that systemic administration of 0.1 mg/kg DCZ did not alter the resting-state functional connectivity. Conversely, 0.3 mg/kg of DCZ was associated with a prominent increase in functional connectivity that was mainly confined to the connections of frontal regions. Additional behavioral tests confirmed a negligible impact of 0.1 mg/kg DCZ on socio-emotional behaviors as well as on reaction time in a probabilistic learning task. 0.3 mg/kg DCZ did, however, slow responses in the probabilistic learning task, suggesting attentional or motivational deficits associated with hyperconnectivity in fronto-temporo-parietal networks. Our study highlights both the excellent selectivity of DCZ as a DREADD actuator, and the side-effects of its excess dosage. The results demonstrate the translational value of rs-fMRI as a drug-screening tool to accelerate the development of chemogenetics in primates.

## INTRODUCTION

The development of chemogenetics such as Designer Receptors Exclusively Activated by Designer Drugs (DREADDs) has ushered in a new era of neuroscience research. DREADDs utilize a family of modified G-protein coupled muscarinic cholinergic receptors to modulate neural activity *in vivo* [1]. In rodents, DREADDs have been instrumental for identifying the function of specific cell populations and neural circuits [2,3]. The development of DREADDs in primates has been slower for a number of reasons [4], but holds the promise of enabling the translation of chemogenetic methods to treat psychiatric and neurological disorders.

DREADD receptors alter the membrane potential of neurons only when bound by non-endogenous ligands [2]. Recently, deschloroclozapine (DCZ) has been identified as a selective and high-affinity activator for muscarinic-based DREADDs with few off-target effects, unlike the original actuator clozapine-N-oxide [5–7]. At a dose of 0.1mg/kg, DCZ occupies up to 80% of DREADD receptors *in vivo* [5], producing rapid and robust changes in neural activity and alterations in behavior. Despite this, there are open questions that could limit the appropriateness of DCZ for use in both basic and clinical settings.

First, it is unclear how DCZ, in the absence of DREADDs, impacts brain-wide patterns of neural activity. This is potentially vital for discerning the more subtle effects of systemic DCZ on the brain, such as changes in functional connectivity between brain areas, information essential for assessing the clinical appropriateness of DCZ. Second, while 0.1 mg/kg DCZ has little impact on tasks that assess executive function or motivation [5,8], it is unknown if the same dose causes changes in translationally relevant affective tasks. While DCZ has negligible affinity for most endogenous receptors, it has some affinity to muscarinic acetylcholine receptors (mAchR) and serotonin type-2 receptors that are primarily located in the cortex [9,10] and which are potentially important for affective behaviors [5]. It is possible therefore that even at low, but functionally relevant doses DCZ may impact affective behavior by binding to these endogenous receptors. Addressing these issues in macaque monkeys is critical as marked changes in brain-wide activity patterns, behavior or both would limit the usefulness of DCZ as the actuator in basic neuroscience and in future clinical settings.

Here we addressed these questions using resting-state functional magnetic resonance imaging (rs-fMRI) and behavioral assessments in macaque monkeys. We measured the impact of either vehicle, low-dose DCZ (0.1 mg/kg), or high-dose DCZ (0.3 mg/kg) on whole-brain functional connectivity as well as in conditioned and unconditioned tasks that assess socio-emotional processing. We found no changes in overall functional connectivity or affective behavior after administration of vehicle or 0.1 mg/kg of DCZ. Conversely, higher doses of DCZ at 0.3 mg/kg affected brain-wide patterns of functional connectivity, in particular in the connections of frontal regions to the temporal and parietal lobes. This dose also impacted decision-making in a probabilistic learning task, increasing reaction times. Thus, our experiments reveal the dosage-dependent impact of DCZ on brain-wide functional connectivity and demonstrate the potential value of rs-fMRI as an additional drug-screening tool in basic and clinical neuroscience.

## METHODS AND MATERIALS

### Subjects

Seven adult rhesus macaques (*Macaca mulatta*) served as subjects (6 males and 1 female). All subjects underwent anesthetized functional MRI scans, while subsets of 4 and 3 animals were assessed in 2 socio-emotional tasks and a probabilistic learning task, respectively (see **Supplemental Table 1**). All procedures were reviewed and approved by the Icahn School of Medicine Animal Care and Use Committee.

### Resting-state fMRI experiments

Animals were first sedated with ketamine (5 mg/kg) and dexmedetomidine (0.0125 mg/kg), and then intubated and maintained under low level (0.7-0.8 %) isoflurane anesthesia [11]. Vehicle (2% DMSO in saline), low-dose DCZ (0.1 mg/kg), or high-dose DCZ (0.3 mg/kg) was administered intravenously halfway through a session. In each session, three to four runs of echo planar image (EPI) functional scans (1.6 mm isotropic, TR/TE 2120/16 ms, flip angle 45°, 300 volumes per run) were obtained for both before (‘baseline’ scans) and after drug injection (‘test’ scans) (**Figure 1**).

**Figure 1.**
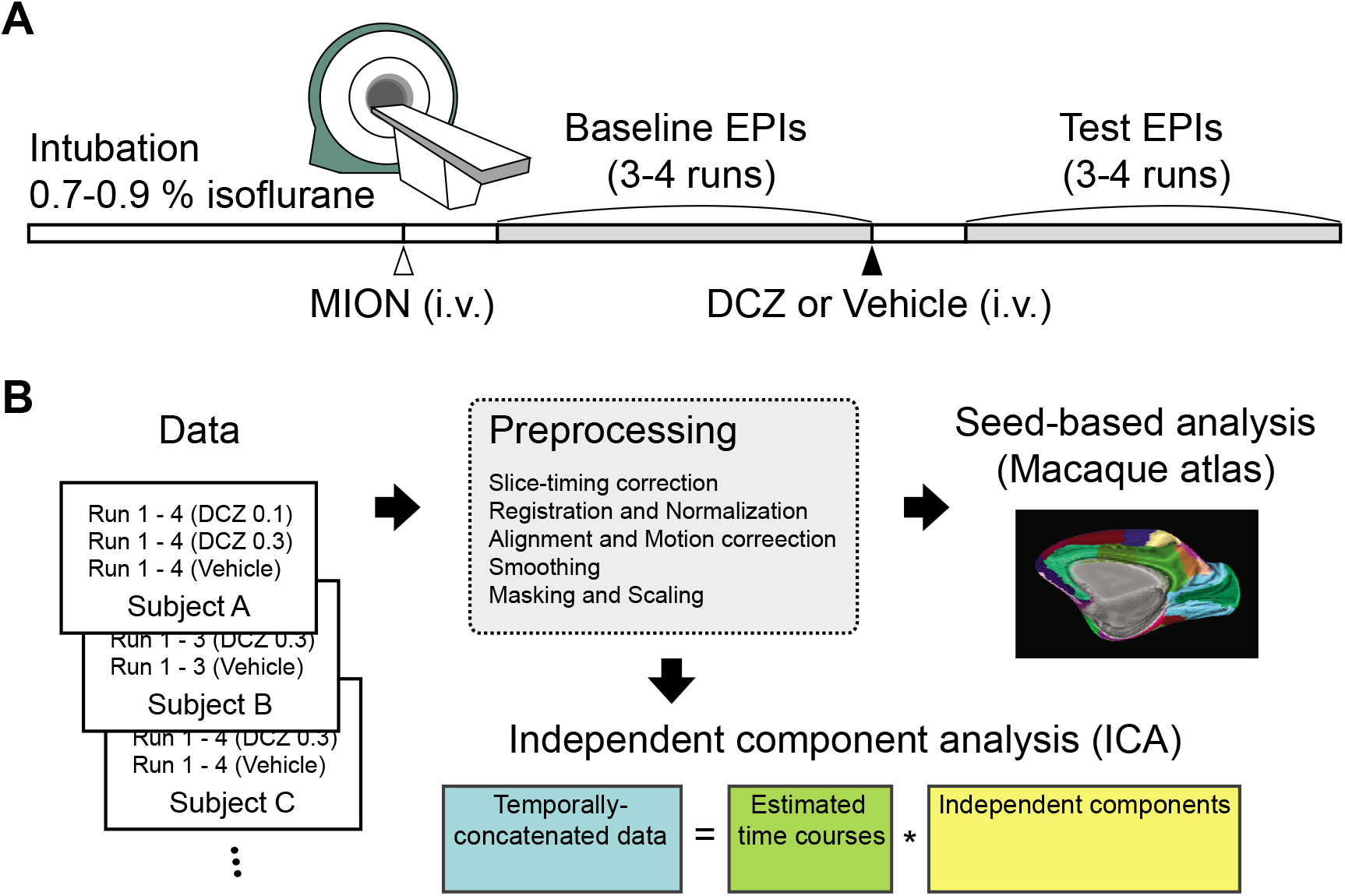
Schematic of experiments. (**A**) An imaging session. Subjects were lightly anesthetized with isoflurane gas throughout a session. A contrast agent (MION, 10 mg/kg) was administered intravenously before functional scans. Drug (vehicle, 0.1 mg/kg DCZ or 0.3 mg/kg DCZ) was prepared and systemically administered (i.v.) following baseline scans, and test scans were acquired after 15 min of drug injection. (**B**) Data analysis pipeline. The scan data for each session initially went through preprocessing procedures. Then seed-based analysis was performed using predetermined ROIs based on the CHARM and SARM atlases. The temporally-concatenated scan data across all sessions was used for ICA.

### Behavioral experiments

Four animals were assessed in the human intruder task [12] and social interaction tests [13]. Vehicle or low-dose DCZ was administered intramuscularly 30-60 minutes prior to a test. Experiments were counterbalanced for treatment order among animals.

Three monkeys were assessed in a probabilistic learning task on a touch-sensitive monitor. In this task, subjects learned to discriminate which visual stimulus in two pairs of novel stimuli was associated with the highest probability (80 or 20%) of receiving a food reward. Each session consisted of 200 trials.

### Data analyses

Connectome analyses were computed using 3dNetCorr function in AFNI [14,15]. We computed the correlation between anatomical ROIs at multiple cortical [16] and subcortical [17] levels in the macaque hierarchical atlases. The correlation matrices, or connectomes, were Fisher’s z-transformed and averaged within condition (baseline and test for: vehicle, low-dose DCZ, and high-dose DCZ). Differences between connectomes were calculated by subtracting the baseline from test data for each drug level. We also calculated a modified group Independent Components Analysis (ICA) using FSL’s melodic ICA function [18,19], with 30 possible components. The extracted IC scores were then submitted to a 2 (baseline or test scan) by 3 (vehicle, low-dose DCZ, or high-dose DCZ) ANOVA which was computed on a voxel-by-voxel basis.

Data from unconditioned socio-emotional tests were scored in Observer XT 14.0 by a condition blind scorer and analyzed using GraphPad Prism 9.0 (GraphPad, San Diego CA). Behavioral ethograms (**Supplemental Tables 2** and **3**) were adapted from previous studies [20,21]. Social interaction was analyzed via paired t-test; human intruder was analyzed via a repeated measures two-way ANOVA with factors of drug and trial type as main factors and monkey as a random effect. The probabilistic learning task was analyzed using MATLAB 2021a. We assessed reaction time (RT, defined as the time from stimuli presentation to a choice response) as a measure of general task-execution processes such as attention or motivation. The effect of drug on RT was assessed using an ANOVA with factors of drug as a main effect and monkey and session as random effects.

Across the rs-fMRI and behavioral testing, we timed injections so that the peak of DCZ concentration in the CSF would occur as testing began [5]. For details of experimental procedures and analyses, see *Supplemental Information*.

## RESULTS

### The impact of low- and high-dose DCZ on brain-wide fMRI functional connectivity

Seven subjects underwent anesthetized functional MRI scanning using isoflurane levels below those shown to adversely impact functional connectivity in frontal cortex [22] (**Figure 1**). To assess the changes in brain-wide functional connectivity following DCZ administration, we calculated functional connectivity using predetermined regions of interest (ROIs) based on the cortical hierarchical atlas (CHARM) [16] and subcortical hierarchical atlas (SARM) [17] for rhesus macaques (**Figure 2A** and **C**). The z-transformed correlation coefficients between fMRI signal time courses in all ROIs were computed for each condition, for both the baseline (before drug injection) and test (after drug injection) scans (**Figure 2B** and **D**, left and center columns). Then, the difference in z-value (test minus baseline) was computed for each drug condition (**Figure 2B** and **D**, right columns). This approach allowed us to control for session-to-session variation in fMRI signals that could influence the results. **Figure 2** shows the full matrix of correlation coefficients from this connectome-based analysis from a single subject (monkey Bu). As expected, we found clear patterns of correlations between distinct brain areas that are consistent to previous reports in macaques [23] in baseline scans (**Figure 2B**, left column).

**Figure 2.**
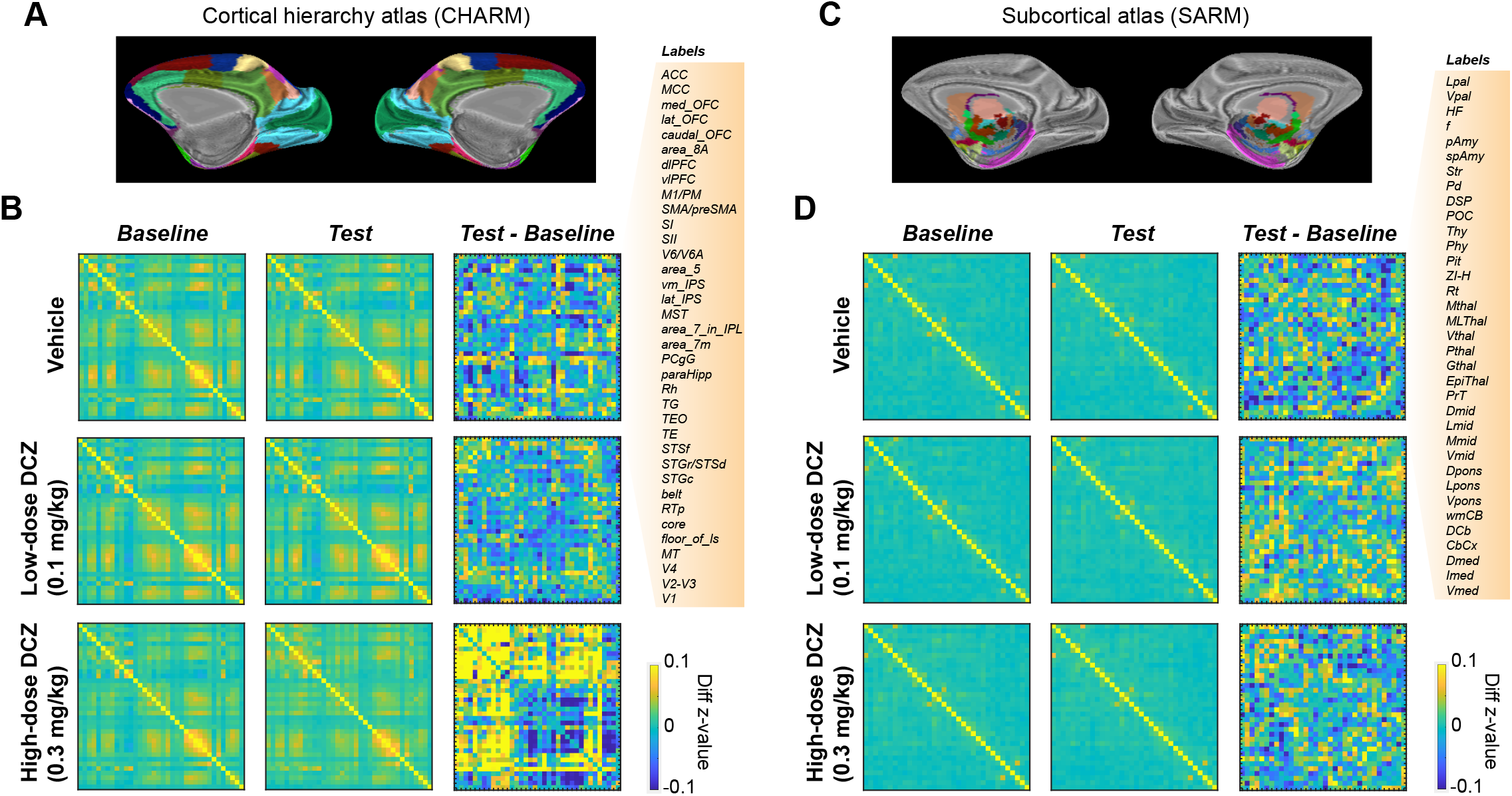
fMRI based connectome changes after DCZ administration in an example subject. (**A**) CHARM atlas level 3 visualized on an inflated brain (medial view). (**B**) Brain-wide functional connectivity in cortical areas. Confusion matrices represent connectome in baseline scans (left column), test scans (center column), and the difference between test and baseline (right column) for either in the vehicle (top), low-dose DCZ (middle), and high-dose DCZ condition (bottom), respectively. Color indicates z-transformed correlation coefficients. Labels on the right side show ROIs from CHARM atlas. (**C**) SARM atlas level 3 visualized on an inflated brain. (**D**) Confusion matrices for subcortical areas with labels for SARM ROIs on the right side. Conventions are as in *B*.

Neither vehicle (2% DMSO in saline) nor low-dose DCZ (0.1 mg/kg) administration caused a detectable change in the overall pattern of functional connectivity and consequently there were very minor differences in functional connectivity between test and baseline scans for these conditions (**Figure 2B**, compare top-right and middle-right panels). Conversely, high-dose DCZ administration (0.3 mg/kg) profoundly increased functional connectivity across cortical regions (**Figure 2B**, bottom-right panel). This effect was most apparent in frontal areas, where, compared to baseline, correlations increased, as well as between frontal cortex and other parts of the brain. In contrast to cortex, correlations between subcortical areas were not impacted by the different drug levels (**Figure 2D**).

To assess the group level impact of DCZ on brain-wide connectomes, we computed correlation matrices for individual subjects for baseline and test scans (N = 5-6 subjects per condition, see **Supplemental Table 1**). Then the z-values differences were averaged across subjects (**Figure 3**). Similar to what we saw for a single subject (**Figure 2**), vehicle or low-dose DCZ had little impact on overall functional connectivity. In contrast, high-dose DCZ increased functional connectivity in cortical areas but not in subcortical areas (**Figure 3A** and **D**). Collectively, average functional connectivity was stronger in the high-dose DCZ condition compared to vehicle or low-dose DCZ in cortical areas but not in subcortical areas (**Figure 3B** and **E**). Two-way ANOVA (Drug: vehicle, low-dose DCZ, or high-dose DCZ × Area category: cortical or subcortical areas) revealed a significant main effect of drug (F_2,13450_ = 31, p = 0.0001) and an interaction between drug and area category (F_2,13450_ = 8.2, p = 0.016). Post-hoc analyses confirmed that high-dose DCZ modulated overall functional connectivity predominantly in cortical areas (vehicle or low-dose DCZ versus high-dose DCZ, p < 0.0001).

**Figure 3.**
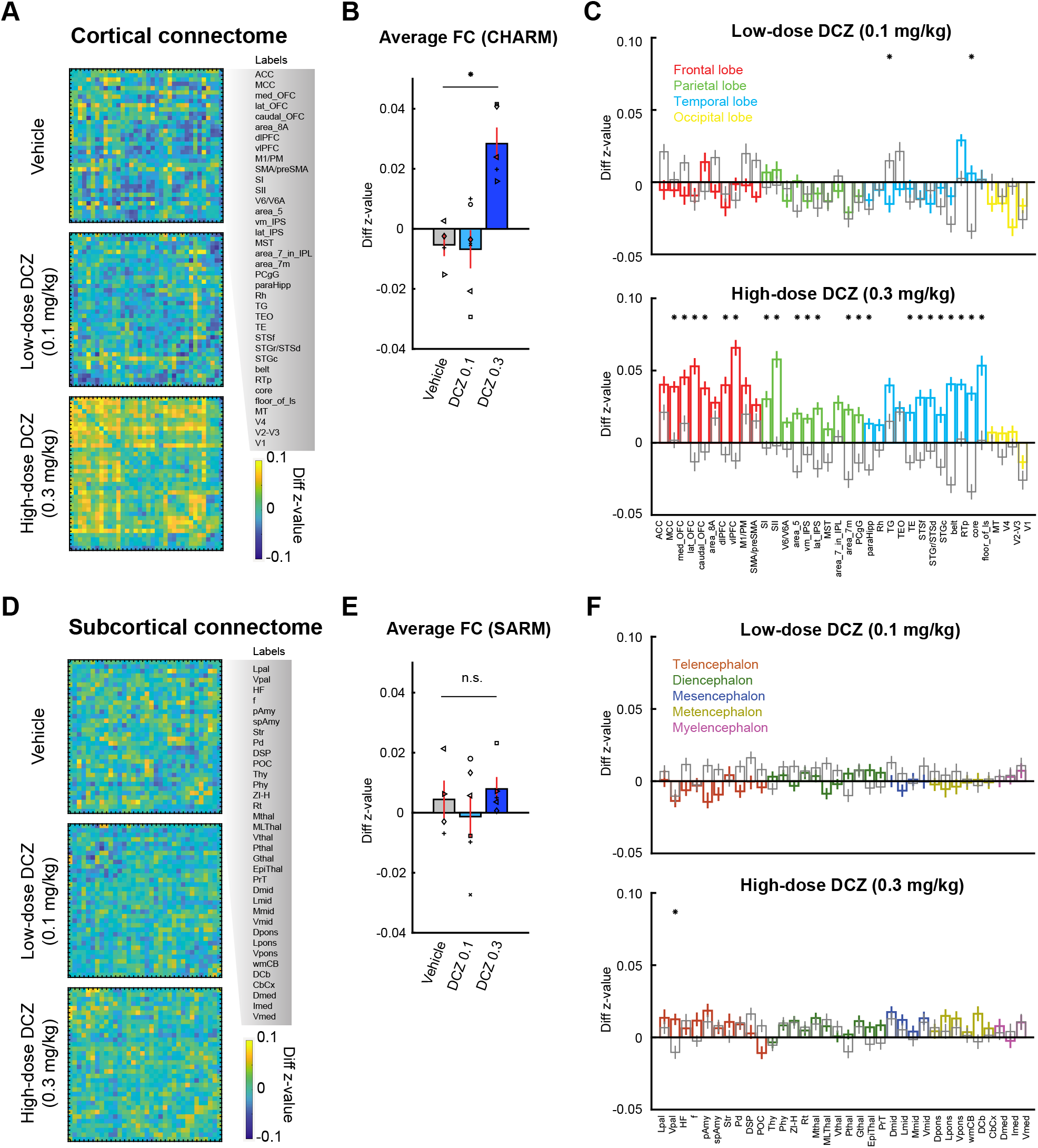
Group level changes in the fMRI-based connectome following DCZ administration. (**A-C**) Functional connectivity changes in cortical areas. (**A**) Confusion matrices represent differences in z-value (test - baseline) for vehicle (top), low-dose DCZ (middle), and high-dose DCZ (bottom) conditions. Labels on the right side show ROIs from CHARM atlas (level 3). (**B**) Averaged functional connectivity. Bars represent averaged z-value differences for vehicle, low-dose DCZ, and high-dose DCZ conditions. Error bars are standard error. Symbols represent subjects. Asterisk indicates a significant interaction of drug by area category (p = 0.016, two-way ANOVA). (**C**) Functional connectivity in each area. Bars indicate averaged z-value differences averaged for each area of the CHARM atlas. Colors indicate frontal (red), parietal (green), temporal (cyan), and occipital areas (yellow) for low-dose (top panel) and high-dose (bottom panel) DCZ conditions. Gray bars indicate vehicle condition. Error bars are standard error. Asterisks indicate significant difference between vehicle and DCZ conditions (p < 0.01 with Bonferroni correction, rank-sum test). (**D-F**) Functional connectivity changes in subcortical areas. Conventions are as in *A-C*. Colors in *F* indicate telencephalon (dark red), diencephalon (dark green), mesencephalon (dark blue), metencephalon (dark yellow), and myelencephalon areas (magenta).

Next, we assessed how high-dose DCZ differentially impacted functional connectivity across different lobes. Frontal, temporal, parietal, but not occipital cortex, showed marked changes in functional connectivity between the high-dose DCZ and vehicle conditions (**Figure 3C**, bottom, * denotes rank-sum test p < 0.01 with Bonferroni correction). Furthermore, the proportion of brain areas that were impacted by low-dose DCZ was far lower than those impacted by high-dose DCZ (**Figure 3C**, top, chi-square test, χ^2^ = 23, p = 1.6 × 10^-6^). Neither DCZ dose level affected functional connectivity in subcortical areas (**Figure 3F**), suggesting the mechanism of functional-connectivity changes was exclusive to cortical regions.

### Independent-components reflect brain-wide functional-connectivity changes caused by high-dose DCZ

While informative, the prior ROI-based approach is limited by predefined anatomical boundaries and may overlook relevant changes in MRI signals not aligned to anatomical boundaries, and does not reveal network-level changes that includes more than two areas. Thus, we performed a data-driven independent-component analysis (ICA) to identify 30 independent components (ICs) that are spatially coherent time courses in the whole-brain fMRI signal. We found a variety of ICs mainly located in cortical areas (**Figure 4A-F**, top). To identify ICs, and thus regions that are affected by high-dose DCZ, a two-way ANOVA was performed (Drug: vehicle, low-dose or high-dose DCZ × Time: baseline or test). This analysis revealed clusters (> 60 voxels with F > 3.5) in a subset of the ICs that showed a significant main effect of drug; these included cortical areas such as the dorso-lateral prefrontal cortex (dlPFC), anterior cingulate cortex (ACC), posterior cingulate cortex (PCC), ventromedial cortex (vmPFC), somatosensory and motor cortices (**Figure 4A-F**, bottom). Notably, IC 3 showed an effect of drug on the fronto-temporo-parietal network, implicated in mediating attentional processes [24] (**Figure 4A**). Further, the ICs which showed drug effects overlapped with regions that showed increased functional connectivity in our ROI-based analyses (**Figure 3**). Systemic administration of high-dose DCZ influenced brain-wide spatially coherent patterns of neural activity, especially in attentionally-linked fronto-temporo-parietal circuits.

**Figure 4.**
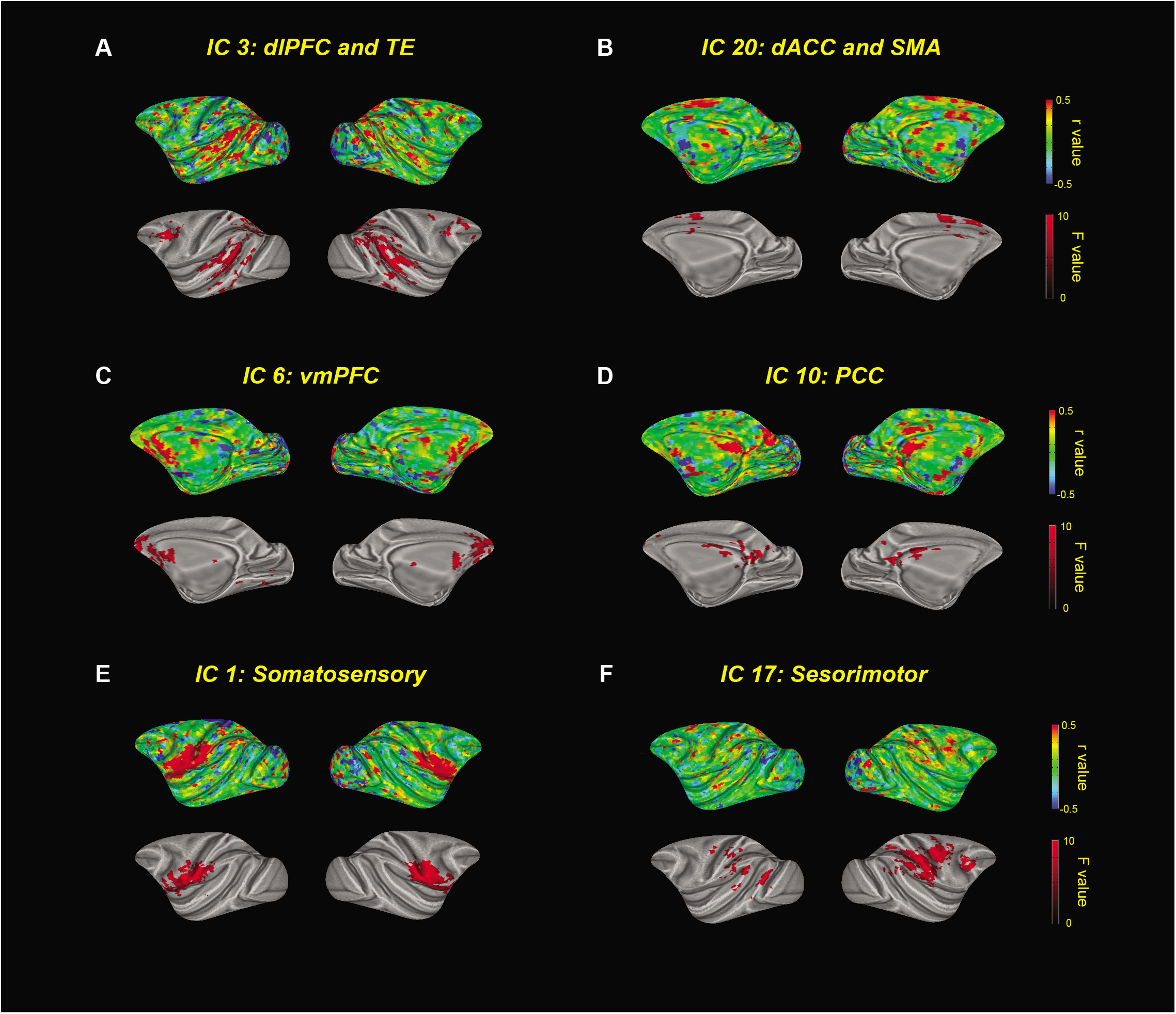
Independent component analysis of the effect of high-dose DCZ. (**A-E**) Top panels show independent components (ICs) as correlation coefficients (*r*) on inflated brains. Bottom panels indicate clusters that showed a significant main effect of drug condition (p < 0.05, two-way ANOVA).

### Negligible effects of low-dose DCZ on socio-emotional behaviors

Our imaging results demonstrated a negligible impact of 0.1 mg/kg DCZ on brain-wide functional connectivity with the least impact in subcortical structures (**Figure 3**). Although no adverse effects of low-dose DCZ were reported in a working-memory task in macaque monkeys [5,8], no study has examined the impact of DCZ on the socio-emotional responses of monkeys. To address this issue, a subset of 4 subjects were assessed in the human intruder task, a well-validated socio-emotional task for non-human primates [12,13]. Subjects received a I.M. injection of either low-dose DCZ or vehicle prior to testing. We analyzed the time spent motionless on each trial type (alone, profile, or stare condition) as a measure of anxiety-like behavior. We expected that the animals spent more time motionless in the ‘profile’ condition in response to indirect threat compared to the ‘alone’ or directly threatening ‘stare’ condition [12]. Animals did spend more time motionless in the ‘profile’ condition in both vehicle and low-dose DCZ conditions (**Figure 5A**). A two-way ANOVA matched for subject (Trial type: alone, profile or stare × Drug: vehicle or low-dose DCZ) revealed a significant main effect of trial type (F_1,3_ = 16, p = 0.049), but no main effect of drug (F_1,3_ = 0.24, p = 0.66) nor an interaction between trial type and drug (F_2,6_ = 1.4, p = 0.32). We also analyzed the frequency of anxious and hostile behaviors during the threatening stare condition. There were no differences between low-dose DCZ and vehicle for either anxiety behaviors (Mean ± SD, vehicle: 10.5 ± 9.6 and low-dose DCZ: 24.8 ± 35.7, respectively; t = 0.68, p = 0.54, paired t-test) or hostile behaviors (Mean ± SD, vehicle: 28.3 ± 32.1 and low-dose DCZ: 37.5 ± 43.4, respectively; t = 1.49, p = 0.23, paired t-test).

**Figure 5.**
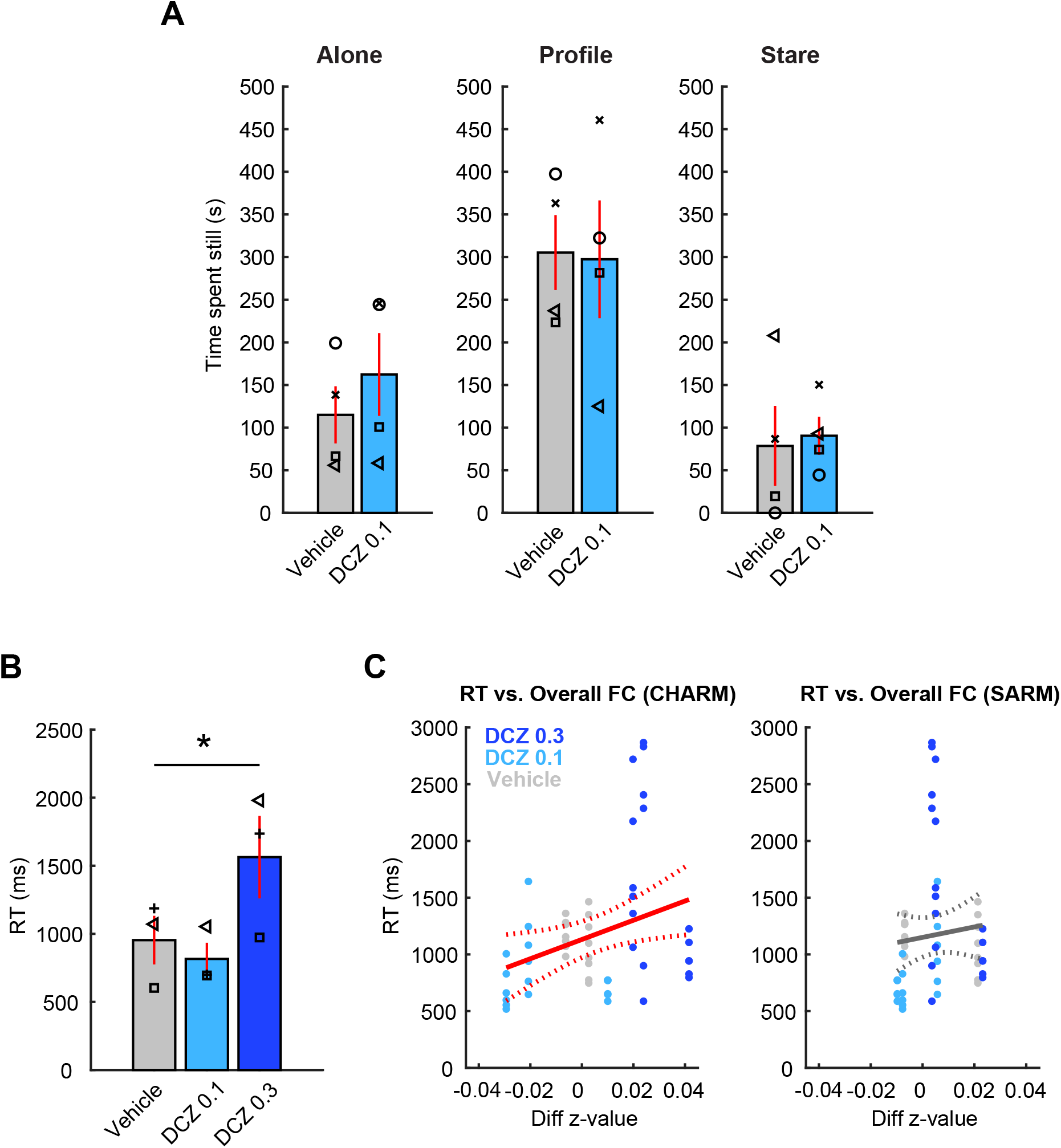
The impact of systemic DCZ administration on socio-emotional behavior and probabilistic learning. (**A**) The effect of low-dose DCZ in the human intruder task. Bars indicate the mean time spent motionless with systemic administration of vehicle (gray) or low-dose DCZ (light blue) either in alone condition (left panel), profile condition (center panel), or stare condition (right panel). Symbols indicate individual animals. Error bars represent standard error. (**B**) Dosage-dependent changes in reaction time (RT) in the probabilistic learning task. Bar plots represent the mean RT of three subjects in each drug condition (vehicle, low-dose DCZ, high-dose DCZ). Error bars are standard error. Symbols represent individual subjects. Asterisk indicates a significant main effect of drug condition (p = 2.5 × 10^-6^, one-way repeated-measures ANOVA). (**C**) Relationship between RT and functional connectivity across conditions in the probabilistic learning task. Scatter plots show the RT in each behavioral session (y axis) and the change in overall functional connectivity in a corresponding subject (x axis) for cortical areas (left panel) and for subcortical areas (right panel), respectively. Individual monkeys completed 6-8 behavioral sessions per each condition. The colors represent drug conditions. The solid and dotted lines are linear fitting and confidence intervals, and red lines indicate significant linear correlation between RT and z-value (p = 0.023).

We also examined whether low-dose DCZ alters social interaction. Subjects were paired with a cagemate, and the total social contact, comprising grooming, play, and passive physical contact, was measured [13]. There were no statistical differences between vehicle and low-dose DCZ conditions on these measures (Mean ± SD, vehicle: 435.4 ± 198.2 s and low-dose DCZ: 502.1 ± 342.3 s, respectively; t = 0.26, p = 0.81, paired t-test). Thus, we found no evidence that macaque socio-emotional behaviors were affected by systemic administration of low-dose DCZ.

### Effect of high-dose DCZ on decision-making and its relation to resting-state functional connectivity

Our imaging results revealed an increase in overall functional connectivity in cortical areas following a systemic administration of high-dose DCZ (0.3 mg/kg) (**Figures 3** and **4**). Because the changes occurred mainly in frontal, temporal and parietal cortex, it likely influences behaviors dependent on functional interaction between these areas (**Figure 4**). We trained a subset of 3 monkeys in a translationally-relevant probabilistic learning task that relies on frontal and temporal cortices [25–27]. We theorized that high-dose DCZ would specifically increase monkeys’ reaction time (RT) as the drug slows decision-making. RTs did increase after high-dose DCZ but not low-dose DCZ, compared to vehicle (**Figure 5B**). A one-way repeated measures ANOVA (Drug: vehicle, low-dose DCZ or high-dose DCZ) revealed a significant main effect of drug (F_2,49_ = 17, p = 2.5 × 10^-6^), suggesting that high-dose DCZ causes an impairment in general task-execution processes, while low-dose DCZ did not. Based on these results, we decided not to include high-dose DCZ in the socioemotional tasks, as an impairment in task execution or attention could compromise the interpretation of the results.

To analyze the relationship between the neural and behavioral effects following systemic DCZ, we correlated overall functional-connectivity changes in cortical (**Figure 5C**, left) or subcortical areas (**Figure 5C,** right) with RT in the 3 animals. There was a significant positive correlation between RT and cortical functional connectivity (n = 58, p = 0.023) but no correlation with subcortical areas (n = 58, p = 0.49), suggesting that cortical resting-state changes following DCZ administration were related to the impairment in task-execution process observed after high-dose DCZ.

## DISCUSSION

To assess the impact of DCZ on brain-wide function in primates, we conducted a resting-state fMRI (rs-fMRI) experiments and behavioral assessments in macaque monkeys. We found that systemic administration of low-dose DCZ, 0.1 mg/kg, did not affect overall functional connectivity or the socio-emotional behaviors we measured. However, we did find that systemic administration of high-dose DCZ, 0.3 mg/kg, increased overall cortical functional connectivity, and was associated with slowed responses in a probabilistic learning task. Together, our data indicate that at a functionally effective dose of 0.1 mg/kg DCZ appears to be appropriate for use as a DREADD actuator in non-human primates, a key finding if chemogenetics are to be translated further.

### Negligible impact of low-dose DCZ on brain-wide functional connectivity or socio-emotional behaviors

Although DCZ was identified as a potent, selective, and metabolically stable DREADD actuator [5], prior work only showed its local effects on the brain using electrophysiology and [18F] FDG-PET imaging [5]. What was not clear is if it impacts brain-wide neural activity in normal controls where DREADD receptors are not present. Consequently, we conducted an rs-fMRI experiment with seven rhesus monkeys and assessed the effect of different dose levels of DCZ. Animals were only lightly anesthetized throughout the scan sessions so that the resting-state brain activity was preserved as closes to that of the awake state as possible [11,22]. We hypothesized that low-dose DCZ, which is functionally effective in chemogenetic experiments in non-human primates [5,28], would not affect resting-state functional connectivity in non-transfected animals. As we expected, low-dose DCZ did not alter brain-wide functional connectivity either in cortical or subcortical areas (**Figures 2** and **3**).

Although DCZ has a high-degree of selectivity to DREADD receptors, it shows some affinity to endogenous muscarinic acetylcholine receptors (mAchR) and serotonin receptors [5]. Past studies demonstrated a negligible effect of low-dose DCZ on performance in a working-memory task in control monkeys [5,7,8], suggesting that this dose of DCZ does not affect working memory performance or attention, processes linked to mAchR [29]. In the current study, we assessed socio-emotional functions of monkeys in the human intruder and social interaction tasks [12,13], which are likely mediated by the serotonergic system [30–33]. We asked whether even subtle agonism to endogenous serotonin receptors by DCZ might impact unconditioned affective behaviors. Our results indicate that systemic low-dose DCZ did not modify affective behaviors of the monkeys (**Figure 5**). In summary, there is a negligible effect of low-dose DCZ on socio-emotional functions or brain-wide functional connectivity.

### Functional-connectivity changes following high-dose DCZ administration

Systemic administration of high-dose, 0.3 mg/kg, DCZ occupies nearly 100% of DREADD receptors when administered in monkeys expressing DREADDs in their brain [5]. The question remained as to whether high-dose DCZ also causes significant non-specific binding of endogenous receptors leading to neural and behavioral alterations. Indeed, DCZ at this dosage reduces performance in a working-memory task, suggesting that the excess DCZ affects neural activity [8]. Our imaging results revealed that high-dose DCZ caused a prominent increase in overall resting-state functional connectivity between frontal, temporal, and parietal lobes that was not observed for vehicle or low-dose DCZ (**Figures 2** and **3**). Notably, the changes were mainly observed in networks that include frontal regions, and no obvious effects were observed in occipital cortex or subcortical structures. This coincides with the pattern of serotonin type-2 receptor (5-HT_2A_, 5-HT_2C_) expression in the brain [34,35], suggesting that excess DCZ may alter brain function through binding to endogenous serotonin receptors.

### Behavioral changes following high-dose DCZ administration were associated with functional connectivity changes

While low-dose DCZ has little impact on behavior, higher dose levels were associated with behavioral changes. Specifically, high-dose DCZ significantly increased RT in the probabilistic learning task compared to other drug conditions, indicating an impairment in general task-execution processes, such as attention (**Figure 5**). Our ICA results revealed a prominent drug effect in a fronto-temporo-parietal network (IC 3, **Figure 4**), heavily linked to attentional processes [24]. Alternatively, high-dose DCZ may increase RT by affecting motivation; this seems less likely, as motivation is largely mediated by subcortical circuits [36].

Past studies have demonstrated that patterns of rs-fMRI functional connectivity in humans are directly related to subsequent behavioral measures recorded in separate sessions [37,38]. Consistently, we found that the changes in cortical functional connectivity and RT were positively correlated (**Figure 5**). This increase in cortical functional connectivity mirrors the slowing of responses may seem counterintuitive as one could expect an improvement in task performance as a result of increased functional connectivity. However, resting-state fMRI functional connectivity reflects low-frequency fluctuations of neural activity between brain regions, and broad, multi-region, increases in functional connectivity do not necessarily benefit coordinated activity across distributed areas [39]. Our data suggests that hyperconnectivity in frontal regions could underlie the impairments in attention that are prominent in neurological and psychiatric disorders [40–42].

### Resting-state MRI as a tool for benchmarking in chemogenetic experiments and drug screening

Primate chemogenetic technique development has been relatively slow for a number of reasons, including the limited number of animals available for development and physiological differences between rodents and primates that hampers viral delivery [4,43]. Although PET imaging is useful for the quantification of chemogenetic receptor expression *in vivo* [5,44–46], the more accessible rs-fMRI should play a complementary role. Here we show that rs-fMRI provides insight into the correlated patterns of neural activity across distinct brain areas and how this can be impacted by pharmacological agents, a level of analysis unavailable with PET. Thus, rs-fMRI has advantages for screening chemogenetic actuators in naive animals without specific task training. Moreover, rs-fMRI is potentially useful for “benchmarking” the selectivity and potency of novel chemogenetic actuators [47], accelerating the development of chemogenetics in primates.

### Conclusion

The present study demonstrated that DCZ is inert in non-DREADD monkeys both in brain-wide activity and in conditioned and unconditioned behaviors, if the dosage is carefully controlled. We also highlight a link between resting-state hyperconnectivity and behavioral impairments, providing a neural mechanism underlying the side-effects caused by excess DCZ. Our data underscore the value of fMRI to accelerate the development of chemogenetics in primates, research that is essential for the translation of these approaches to treat human psychiatric disorders.

## ACKNOWLEDGEMENTS

AF, CE, BER and PHR are supported by grants from NIMH and the BRAIN initiative (R01MH110822 and R01MH117040). BER is supported by grants from NIMH (R01MH111439) and NINDS (R01NS109498). AF is supported by Overseas Research Fellowship from Takeda Science Foundation and a Brain & Behavior Research Foundation Young Investigator grant (#28979). We would like to thank Dr Paula Croxson for providing the foundation on which this work was built and Jairo Munoz and Niranjana Bienkowska for assistance with data acquisition. For help with fMRI data pre-processing and analysis we thank Drs Paul Taylor and Alex Franco, respectively.

## AUTHOR CONTRIBUTIONS

AF, CE, PHR and BER developed the experimental design and ideas for the research; AF, CE, JMF, SHF and LF performed the research; AF, CE and BER analyzed the data; AF, CE, PHR and BER wrote the manuscript; all authors edited the manuscript.

## SUPPLEMENTAL INFORMATION

### SUPPLEMENTAL METHODS

#### Drug preparation

To prepare a DCZ solution for fMRI or socio-emotional testing, DCZ powder (HY-42110, MedChemExpress) was dissolved in dimethyl sulfoxide (DMSO, 2% of total volume) and then diluted in saline to a concentration of 0.1 mg/kg or 0.3 mg/kg, with a total volume of 1ml. Vehicle solution was prepared as 2% DMSO diluted in 1ml saline. One vehicle scan was conducted without DMSO (monkey Wo). Both solutions underwent a sterilized filter (0.22 *μ*m) before injection. DCZ or vehicle solution was prepared within 30 min of usage.

For the probabilistic learning testing, both vehicle and DCZ were prepared using sterile procedures and previously published drug preparation approaches for behavioral testing in macaques [1]. The vehicle consisted of 0.5% Acetic acid, 50% 1 M Sodium Acetate, and 49.5% 0.2 M NaOH. Each solution of DCZ or vehicle ultimately had a pH between 5.5 and 6.0. When DCZ was administered, the drug was first dissolved in acetic acid and sodium acetate and then diluted with NaOH. The vehicle remained consistent across both drug doses (0.1mg/kg, or 0.3mg/kg of DCZ). However, the concentration of each dose was altered so the total volume of each injection was the same no matter the dose, e.g. a 0.1mg/kg dose had a concentration of 1 mg/mL, while a 0.3 mg/kg dose had a concentration of 3 mg/mL. As with fMRI testing, DCZ or vehicle solution was prepared within 30 min of usage.

#### fMRI data acquisition

Animals were first sedated with ketamine (5 mg/kg) and dexmedetomidine (0.0125 mg/kg) at least 1.5 hours before the data collection to prevent detrimental effects of ketamine on neural activities. They were then intubated and maintained under low level (0.7-0.8 %) isoflurane throughout the session to ensure preservation of resting-state networks [2]. During all scanning sessions, anesthetized macaques were contained within a primate chair in the sphinx position with their heads supported by a sling. Because animals were not held in a stereotactic frame, this meant we were able to maintain sedation with lower levels of anesthesia than is often used. To minimize the effect of physiological changes in the neural activity, vital signs (end-tidal CO2, body temperature, blood pressure, capnograph) were continuously monitored and maintained as steadily as possible throughout an experimental session. Structural and functional scans were collected each session, as well as pre-injection (baseline) and post-injection (DCZ or vehicle) functional scans. First, a set of setup scans were acquired which included shimming based off the acquired fieldmap. Additionally, a 3D T1-weighted image (0.5 mm isotropic, TR/TE 2500/2.81 ms, flip angle 8°) was acquired. Following MION injection (i.v.) [3,4], three to four runs of echo planar image (EPI) functional scans (1.6 mm isotropic, TR/TE 2120/16 ms, flip angle 45°, 300 volumes per each run) were obtained for baseline functional resting states. Then, approximately 30 minutes from the beginning of the pre-injection rs-fMRI data collection, either DCZ (0.1 or 0.3 mg/kg) or vehicle was administered (i.v.), and subsequently an additional three to four functional runs (with the same parameters) were collected. Test EPI data were collected approximately 15 minutes after DCZ or vehicle injection to ensure trafficking of the drug to the brain [5].

#### Unconditioned socio-emotional tasks

Four animals (Wo, He, La, Ha) were tested on two well validated behavioral tasks that assess socio-emotional function: the human intruder [6] and social interaction tests [7]. Behavioral tests were conducted following injections of either vehicle (2% DMSO in saline) or 0.1 mg/kg DCZ. Injections were given intramuscularly (i.m.) 30-60 minutes before behavioral testing in keeping with the known pharmacokinetics of DCZ (Nagai et al. 2020). Three of the animals received vehicle treatments before low dose DCZ treatments, and a fourth animal received low dose DCZ treatment before vehicle. Each animal was tested only once in each drug condition for each behavioral test. All behavioral data were scored offline in Observer XT version 14.0 (Noldus, Wageningen, the Netherlands).

##### Human Intruder

Animals were tested on a human intruder paradigm, which has been shown to accurately assess anxiety behaviors in macaques [6,8–10]. Testing was conducted in a Wisconsin General Testing Apparatus (WGTA). The sliding doors of the WGTA were left open so the animal could observe the room throughout the test session. We have adapted our testing procedure and behavioral ethogram from a previous study [11].

The paradigm consisted of three conditions (alone, profile, stare) presented in the same order to all monkeys for a total duration of 30 min. The experimenter wore a rubber mask depicting a different human male face during each testing session, presented in pseudorandom order to subjects with no repetitions of masks per subject. Different colored curtains hung behind the experimenter’s seat were also used to increase novelty. Within each testing session, monkeys were first left alone to observe the room and acclimatize to the environment (alone condition). Next, the experimenter, wearing a novel mask, stepped into the animal’s field of view with only their profile visible to the animal, then sat in a chair at the eye level of the animal for the duration of the condition. The experimenter remained still, with only the profile visible, for 9 minutes (profile condition). Another alone condition elapsed for 3 minutes in order to allow the animal to return to baseline behavior, before the experimenter stepped back into view, then sat in a chair at eye level facing the animal and making eye contact. The experimenter remained still, attempting to make eye contact with the animal for 9 minutes (stare condition) before stepping out of view and ending the session. A minimum of one month elapsed between repetitions of this task with the same animal, as animals may habituate to the human intruder (Kalin et al. 2018).

##### Social Interaction

Animals were removed from the home cage and placed in a play cage set up in an unfamiliar room. A familiar cagemate was introduced to the play cage, and social interaction between the subject and cagemate was observed. We used an ethogram previously defined by others to evaluate social behaviors [12,13]. The cagemate chosen for this task was always the same across conditions. The cagemate did not receive any injection prior to testing. A minimum of 48 hours elapsed between social interaction tests.

#### Probabilistic stimulus-reward learning task

Three monkeys (He, Cy, and Wo) were trained to perform a two alternative forced choice probabilistic learning task on a touch-sensitive monitor. All testing occurred while monkeys were seated in a wheeled transport cage. In each 200-trial session, subjects had to learn to discriminate which visual stimulus in two pairs of novel stimuli was associated with the highest probability of receiving a food reward pellet (banana, grape, and raspberry flavored 20mg BIO-Serv Dustless Precision Pellets^®^). In each pair of stimuli, one option was associated with an 80% probability of receiving a reward whereas the other was associated with 20% probability of receiving a reward. Stimuli were novel at the beginning of each session, the assignment of visual stimuli to high/low probability was random, and the probability of receiving a reward was fixed throughout the session.

Each trial began with the presentation of a centrally located green square ‘lever’ stimulus. Once monkeys pressed the lever stimulus, two stimuli were presented to the left and right of the screen, and monkeys could choose between them. Pressing one stimulus caused the non-selected option to be removed from the screen. The chosen stimulus remained on the screen for a further 0.5 seconds before being extinguished and a 4 second inter-trial interval commenced. For one pair of stimuli, if a reward was to be delivered it was dispensed when the chosen stimulus was removed from the screen. For the other pair of stimuli if a reward was to be delivered, it was dispensed 1 second after the chosen stimulus was removed from the screen. The presentation of stimuli pairs was randomized across trials and the left-right location of each stimulus was also counterbalanced across trials.

Monkeys completed daily 200-trial sessions (100 trials of each pair), run 4-5 days per week. 0.1mg/kg, 0.3mg/kg DCZ, and vehicle were administered in sessions after stable performance was established each week. In practice this meant that injections of 0.1mg/kg, 0.3mg/kg DCZ, or vehicle were administered on either Wednesday, Thursday, or Friday.

#### fMRI analysis

Functional imaging data was preprocessed with standard AFNI/SUMA pipelines [14,15]. Raw images were first converted into NIFTI data file format and ordered into BIDS format [16]. The T1 weighted image from each session was first warped to the standard NMT atlas space using a newly developed macaque skull stripping function followed by AFNI’s @animal_warper command [15,17]. Specifically, the T1w images were spatially normalized, then skull stripped using the U-Net model built from Primate Data-Exchange open data sets [18], the mask was dilated out 6mm to ensure the whole brain was within the mask, and finally the mask was applied to the original image, which was then renormalized. This normalized skull-stripped T1w image was then aligned to the NMT atlas [17], along with subject specific versions of the CHARM [15] and SARM atlases [19].

The EPI data were preprocessed using a customized version of the AFNI NHP preprocessing pipeline (Jung et al 2020). For each session pre- and post-injection data were processed separately using the same parameters. In addition to a set of dummy scans, the first 3 TRs of each EPI were removed to ensure that any magnetization effects were removed from the data prior to functional connectivity analyses. The images were first slice time corrected, then motion correction was applied and the EPIs were aligned to the within session T1w image and warped to the standard space. Following alignment to the standard space, the EPIs were blurred with a full width half max of 3mm, and then converted to percent signal change. We then regressed the motion derivatives from each scan along with CSF and WM signal regressors. The residuals of this analysis were then used to compute the functional connectivity analysis described below.

The full connectome analyses were computed using 3dNetCorr function in AFNI [14,20]. The error time series from the preprocessing steps described above were split into half runs and then we computed the correlation between ROIs at various levels of the cortical [15] and subcortical [19] hierarchical atlas. Following the creation of the individual correlation matrices, or connectomes, for each half run, the connectomes were Fisher’s z-transformed and then averaged within condition to create mean connectomes for each condition (both pre-/post-injection for the vehicle, DCZ 0.1 mg/kg, and DCZ 0.3 mg/kg sessions) at all hierarchical levels. We then calculated the difference between connectomes by subtracting the pre-injection data from the post-injection data for each session. Finally the average difference connectome was computed across subjects for each drug level. To statistically determine the effect of drug on the functional connectivity in CHARM and SARM atlases, the mean connectomes from individual monkeys were then submitted to an ANOVA with main effects of drug and area category. Monkey and brain area were modeled as random effects and brain area was nested under monkey. In addition to the functional connectomes analyses, we used a modified group Independent Components Analysis (ICA) to look for any changes in the component structure of our data following the drug manipulations. The same split run error time series from the connectome analyses were submitted to FSL’s melodic ICA function [21,22] allowing for 30 possible components and a mixture model threshold of 0.5. Following the computation of the group ICA, we submitted resultant data to FSL’s dual regression function, which was used to extract the spatial ICs for each subject and condition [21,23]. The extracted condition IC scores were then submitted to a 2 (Pre-Injection vs Post-Injection) by 3 (Drug: Vehicle, DCZ 0.1 mg/kg, DCZ 0.3 mg/kg) ANOVA which was computed on a voxel by voxel basis.

#### Behavioral analysis

Data from socio-emotional tests were assessed by a scorer blinded to the condition. Data was scored in Observer XT 14.0 and exported to GraphPad Prism 9.0 (GraphPad, San Diego CA). Behavioral ethograms (**Supplemental Tables 2** and **3**) were adapted from previous studies [11,13]. Behavioral responses in the human intruder and social interaction paradigms were assessed for normality using a Shapiro-Wilk normality test. Human intruder data was analyzed using two-way repeated measures ANOVAs with matching by subject. No assumption of sphericity was made. Sidak’s multiple comparison test was used to evaluate differences between categories (i.e. differences between DCZ and vehicle treatments within the alone, profile, and stare conditions). Hostile and anxious behaviors during the stare condition of the human intruder test were analyzed using a paired student’s t-test, with matching by subject. Social interaction data was also analyzed using a paired student’s t-test, with matching by subject.

Data from the probabilistic learning task was analyzed using MATLAB 2021a. In all sessions analyzed, each of the monkeys learned to discriminate the pairs of stimuli by the end of the session. This was based on a running average of 10 trials and criterion was set at 8/10 trials or 80%. We assessed reaction time (RT) as a measure of general task-execution process such as attention or motivation. RT was defined as the time duration (in milliseconds) from the presentation of two stimuli to a choice response of the monkeys. We analyzed RT using ANOVA with factors of drug as a main effect and monkey and session as random effects.

## SUPPLEMENTAL TABLES

**Supplemental Table 1:** Assignments of monkeys to fMRI and behavioral testing conditions. Y and N indicate the condition that the data was collected and not collected, respectively.

| Animal | rs-fMRI<br>(vehicle) | rs-fMRI (DCZ 0.1<br>mg/kg) | rs-fMRI (DCZ 0.3<br>mg/kg) | Socio-emotional<br>tasks | Probabilistic learning<br>task |
| --- | --- | --- | --- | --- | --- |
| La | N | Y | N | Y | N |
| Ha | Y | Y | N | Y | N |
| Wo | Y | Y | Y | Y | N |
| He | N | Y | Y | Y | Y |
| Cy | Y | Y | Y | N | Y |
| Bu | Y | Y | Y | N | Y |
| Pi | Y | N | Y | N | N |

**Supplemental Table 2.** Behavioral ethogram for human intruder task, adapted from Raper et al. 2019.

| Category and Specific Behaviors | Measurement | Examples |
| --- | --- | --- |
| Affiliative | Cumulative Frequency |  |
| Lipsmack | Frequency | Rapid movement of pursed lips, accompanied by a smacking sound |
| Present | Frequency | Rigid posture (knees locked) with tail elevated and rump oriented toward the stimulus object |
| Anxiety | Cumulative Frequency |  |
| Scratch | Frequency | Rapid scratch of body with hands or feet |
| Body shake | Frequency | Shake of the whole body or just head and shoulders region |
| Tooth grind | Frequency | Repetitive rubbing or chewing of upper and lower teeth |
| Yawn | Frequency | Open mouth widely, exposing teeth |
| Fearful | Cumulative Frequency |  |
| Withdrawal | Frequency | Quick, jerky motion away from the stimulus object (jump back) |
| Grimace | Frequency | Retracted lips, exposed clenched teeth |
| Hostile | Cumulative Frequency |  |
| Cage aggression | Frequency | Slaps, shakes, bites, or slams body against cage |
| Threat | Frequency | Open mouth (no teeth exposed) |
| Bark | Frequency | Low pitch, high intensity, rasping, guttural |
| Lunge | Frequency | A quick, jerky movement toward the stimulus |
| Position | Duration |  |
| Front of cage | Duration | No part of animal's body contacts back of the cage |
| Back of cage | Duration | Animal's body contacts back of the cage |
| Stereotypies | Cumulative Duration |  |
| Pacing | Duration | Repetitive motor pattern around the test cage |
| Motor Stereotypy | Duration | Repetitive, abnormal voluntary or involuntary motor patterns (e.g. swinging, twirling, flipping) |
| Stillness | Duration | Motionless posture for at least 3 seconds (except small head movements) |
| Vocalizations | Cumulative Frequency |  |
| Coo | Frequency | Clear soft pitch and intensity, sounds like “ooooh” |
| Scream | Frequency | High pitch, high intensity screech or loud chirp |

**Supplemental Table 3.**
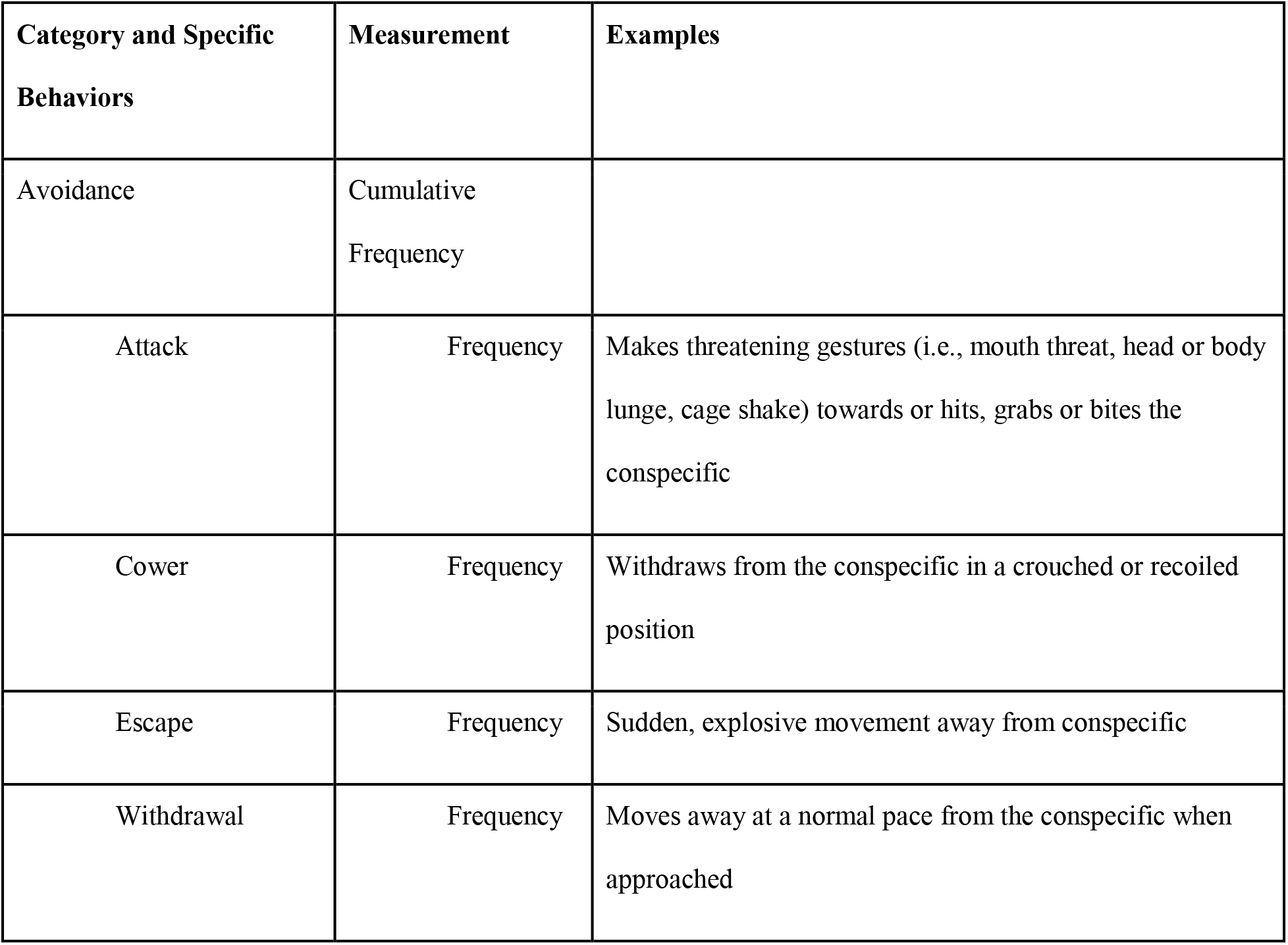
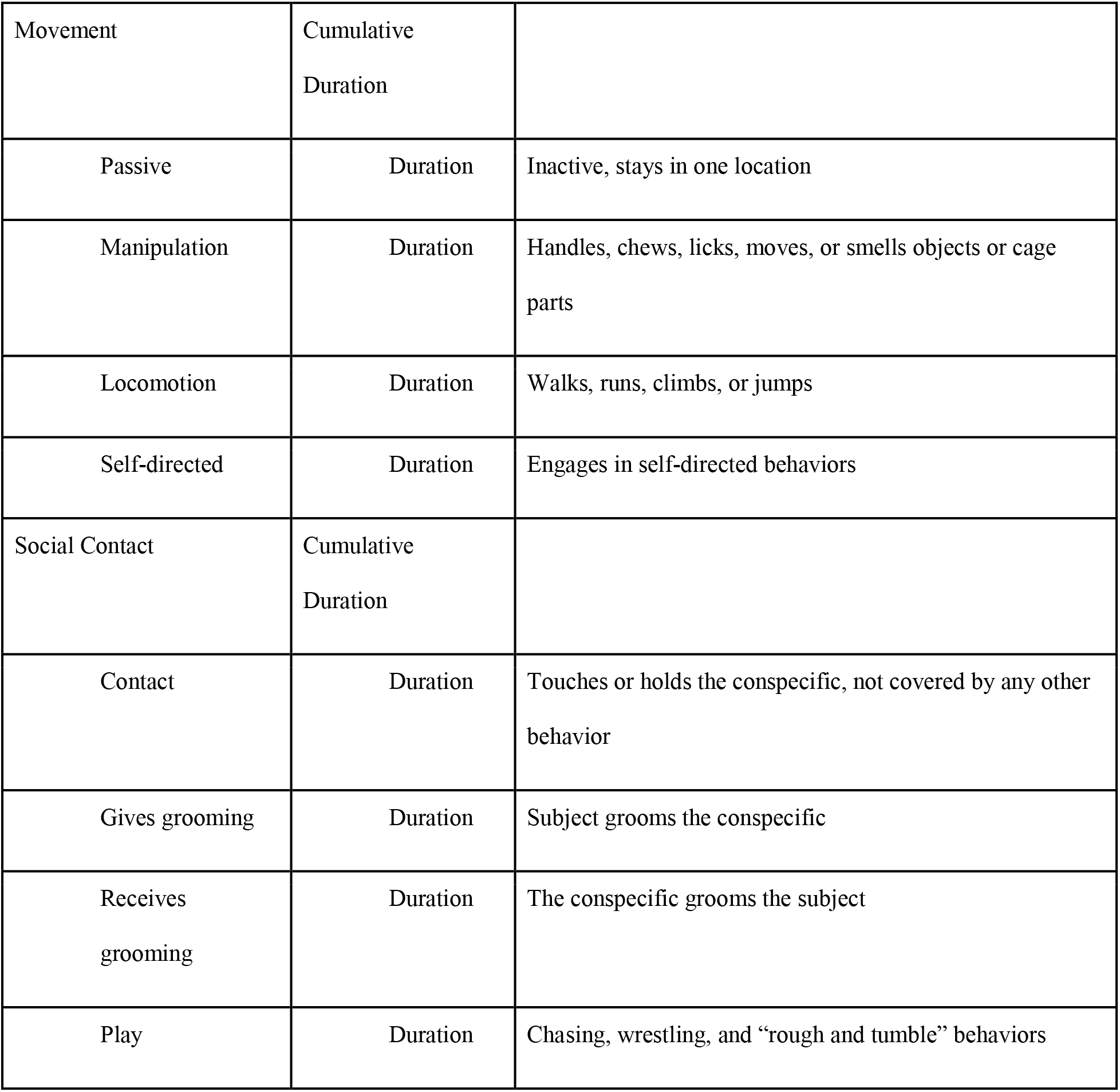
Behavioral ethogram for social interaction task, adapted from Forcelli et al. 2017.

| Category and Specific Behaviors | Measurement | Examples |
| --- | --- | --- |
| Avoidance | Cumulative Frequency |  |
| Attack | Frequency | Makes threatening gestures (i.e., mouth threat, head or body lunge, cage shake) towards or hits, grabs or bites the conspecific |
| Cower | Frequency | Withdraws from the conspecific in a crouched or recoiled position |
| Escape | Frequency | Sudden, explosive movement away from conspecific |
| Withdrawal | Frequency | Moves away at a normal pace from the conspecific when approached |
| Movement | Cumulative<br>Duration |  |
| Passive | Duration | Inactive, stays in one location |
| Manipulation | Duration | Handles, chews, licks, moves, or smells objects or cage parts |
| Locomotion | Duration | Walks, runs, climbs, or jumps |
| Self-directed | Duration | Engages in self-directed behaviors |
| Social Contact | Cumulative<br>Duration |  |
| Contact | Duration | Touches or holds the conspecific, not covered by any other behavior |
| Gives grooming | Duration | Subject grooms the conspecific |
| Receives grooming | Duration | The conspecific grooms the subject |
| Play | Duration | Chasing, wrestling, and “rough and tumble” behaviors |

